# Extending Conway’s Game of Life

**DOI:** 10.1101/2022.08.30.505937

**Authors:** Wayne M Getz, Richard Salter

## Abstract

We introduce an extension to Conway’s Game of Life (GoL) that allows more than 2 states (dead/alive; 0/1) to be specified. State 0 is reserved for dead or lifeless cells, and states 1 to *n* − 1 for cells that are “alive.” Associated with state *σ* is the value *v*_*σ*_ of cells such that *v*_0_ = 0, *v*_1_ = 1, and for *σ* = 2, …, *n* − 1 the cell value *v*_*σ*_ is free to be assigned any positive integer value. The value *V*_*ij*_ of the Moore neighborhood surrounding cells at array locations (*i, j*) (*i* = 1, …, *r* rows, *j* = 1, …, *c*) is the sum of state values of the 8 cells that touch the cell at location (*i, j*) (4 north, south, east and west side cells and 4 NE, NW, SE, and SW corner cells; except for boundary cells, which have 5 neighbors on an edge or 3 neighbors on a corner, and noting that torroidal arrays have no boundaries). The game is based on a set of rules involving the values *V*_*ij*_ that determine whether cells at locations (*i, j*) in state *σ*, stay in this state or progress to state *σ* + 1, *σ* = 0, …, *n* − 2, or die (remain dead in the case of dead cells). Specifically, for two non-negative constants 0 ≤ *p*_*σL*_ ≤ *p*_*σU*_, if *V*_*ij*_ *< p*_*σL*_, *V*_*ij*_ ∈ [*p*_*σL*_, *p*_*σU*_] or *V*_*ij*_ *> p*_*σU*_ then a cell in state *σ* at location (*i, j*) respectively stays in state *σ*, progresses to state *σ* + 1, or dies. In this paper, we illustrate some of the properties of such games of life with three, four and six states, identifying still-life, oscillator, glider and replicator pattern sequences. We also examine the long term behavior of patterns arising from random and regular (stripped diagonals) starting configurations, as well as set-piece motifs. Most importantly, we introduce two freely downloadable, Numerus runtime alterable model platforms (Numerus RAMPs) that can be used to simulate the three and four state GoLs discussed in this paper, and provide a guide on how these RAMPs can be used to explore the behavior of our GoLs.

## Introduction

Conway’s “Game of Life” (GoL) was first introduced to a general scientific audience by Martin Gardner in his October 1970 Scientific American column Mathematical Games (Gardner, 1970). This game captured the imagination of mathematical, physical, and biological scientists, as well as the scientifically curious lay public. This game provides a startling example of how an extremely simple, two-dimensional cellular automaton exhibits complex dynamical systems properties. These properties include chaotic and self-organizing behaviors, phase transitions, self-replicating pattern generation, and self-organized criticality (Alstrøm & Leão, 1994; Caballero et al., 2016). Notably, the game of life itself has been shown by computer scientists to be a Universal Turning Machine in being able to carry out, albeit in an extremely laborious fashion, any computation that a modern general purpose electronic computer can perform (Berlekamp et al., 2004).

Conway’s GoL also provides a wonderful tool for capturing the imagination of school kids, even at the elementary school level. It stimulates them to experiment with starting configurations that produce unexpected patterns of behavior as time progresses. Conway’s GoL has become so popular, that a Google search in June, 2022, using the phrase “Conway’s game of life simulator” produced 66,500 hits. The top hit was `playgameoflife`. This site provides a highly versatile GoL simulation platform that is implemented in the users browser.

The GoL is played out on cellular array consisting of a user-selected number of rows and columns (essentially a large rectangular checkered board where at the start of the simulation any cell may be have one of two colors—e.g. grey and yellow, as in Fig. 1). At the `playgameoflif` site, the user is able to choose among thousands of named starting configurations (such `ants, big glider, super fountain, toad`, and `zweiback`), control the speed of the simulation, move through the simulation one step at a time, alter the starting configuration (aka initial conditions) by clicking the mouse on individual squares and thereby changing its current color, and setting the size of the cellular array on which the simulation plays out. It also provides a graphical illustration of the GoL rules from cells to switch from being dead to live and vice-versa (Fig. 1)

**Figure 1.**
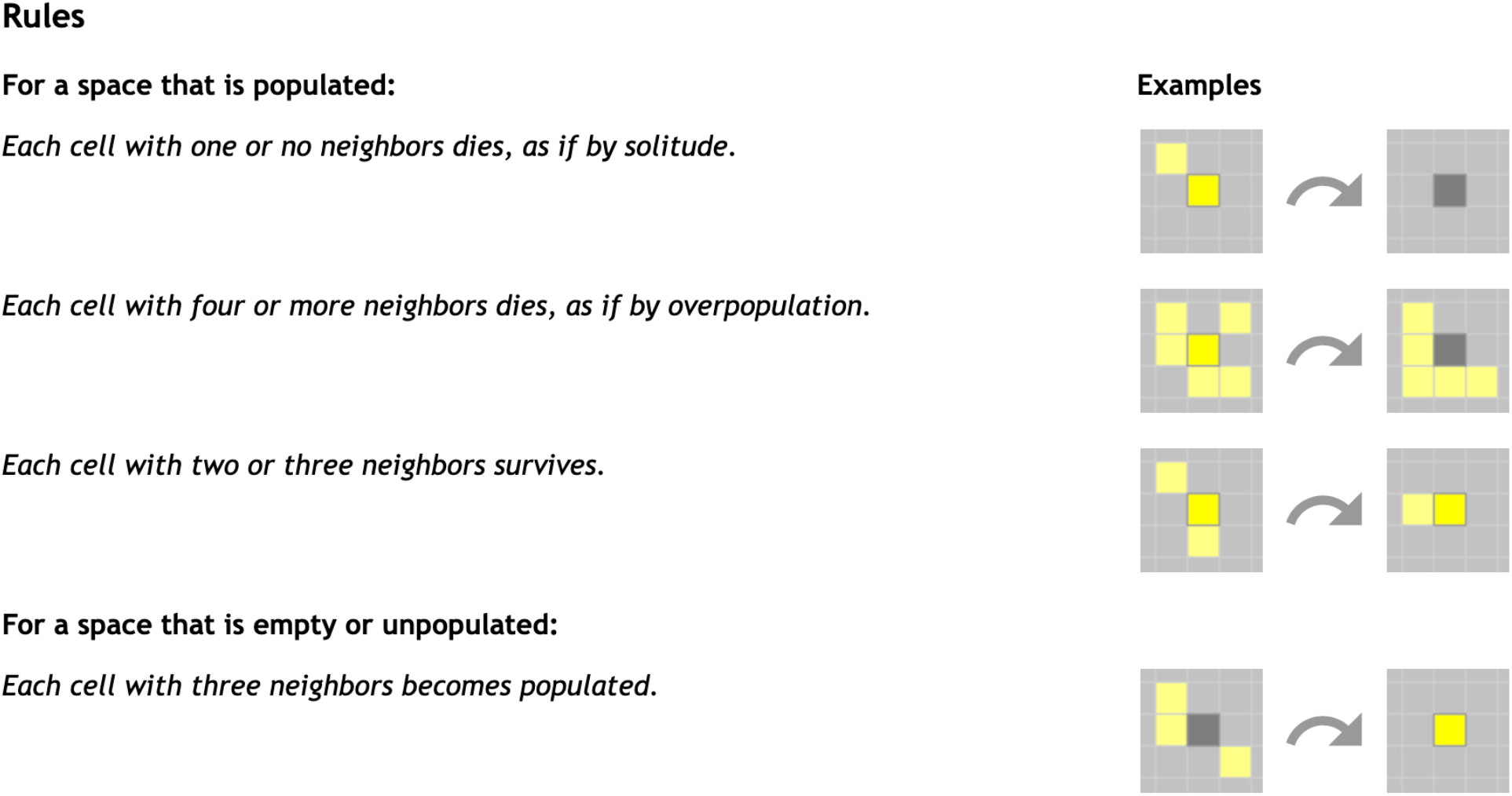
A graphical depiction of Conway’s Game of Life (GoL) played out on a rectangular grid of cells (aka an array). Yellow cells are “live” (state value 1) and gray cells are “dead” (state value 0). The focal or centered cell referred to in the text is highlighted in a more solid shade of yellow or gray, as the case may be, than the cells in its Moore neighborhood of eight surrounding cells. This figure has been cut out from a webpage that is part of the `playgameoflife` online implementation.

The GoL has now been around for more than 5 decades, so “all the low hanging fruit” discoveries have been made. Obtaining new results and insights now requires some work. In contrast, the extended versions of the GoL that we present here have “loads of low hanging fruit.” These versions thus provide opportunities for students to experiment, explore, learn and discover new constructs and phenomena without the need for deep mathematical insights into the complex systems behavior of automata. Further, we have implemented some of these extended GoLs as Numerus^®^ Runtime Alterable Model Platforms (RAMP). The platforms employs a technology that allow the user to replace or build small segments of code called RAMs (runtime alternative modules) on the fly (i.e., while in the runtime environment). Our RAMPs and RAMP player, `Numerus Studio`^®^, are free, simple to use, and run under MacOS, or Windows operating systems.

In the remainder of this paper we present the rules of the game, demonstrate the behavior of several versions of the game, and provide a guide to the use of our RAMPs in the classroom.

### Rectangular Arrays and Conway’s Game of Life

#### Definitions

For the sake of completeness, we first provide a very brief review of rectangular array neighborhoods and the rules of Conway’s GoL. We first note the GoL is played on a two-dimensional rectangular grid containing *r* rows and *c* columns of cells. It is typically up to the user to choose values for *r* and *c*, with 30 to 100 being a common range for each of these two parameters.

Apart from boundary cells in the rectangular array, each internal cell has four neighbors on its four sides and an additional four neighbors at its four corners. In cellular array nomenclature, the four side-abutting neighbors are said to constitute the *von Neumann neighborhood* of each focal cell. These four von Neumann neighborhood cells with the addition of the four corner-abutting neighbors is said to consitute the *Moore neighborhood* of each focal cell. It now remains to define the neighborhoods of boundary cells. Boundaries can either be an edge (rectangular arrays) or a wrap (toroidal arrays). Cells on edge boundaries have fewer neighbors. For Moore neighborhoods, edge boundary corner cells have only three rather than eight neighbors and each edge boundary non-corner cell has five cells in its Moore neighborhood. Wrapped boundary cells have their full complement of neighborhood cells, because the north and south boundary cells, as well as the east and west boundary cells are now neighbors. From a topological point of view, linking the north and south boundaries turns our rectangular array into a horizontal cylindrical array, and then linking the two ends of this cylinder turns the array into a torus or toroidal array (colloquially referred to as a doughnut).

Conways GoL is played out on an *r* × *c* arrays that either has edge (rectangular array) or wrapped (toroidal array) boundaries. Each of the cells in these two types of arrays can have one of two states: 0 (aka dead) and 1 (aka live). For purpose of visual presentation, a color is generally associated with each state (typically white or gray with 0) and (typically black or some vibrant color with 1). At the start of the game, also referred to as the simulation, each cell is assigned an initial state value. This assignment can be coded to be random or regular (e.g., some type of stripped pattern), or by clicking a mouse on a cell to change its state from 0 to 1 (e.g., the gameoflife website implementation). At each new step of the simulation—aka tick of the time clock *t* = 0, 1, 2, 3, …, —the current state of each cell at time *t* is updated to its new or next state at time *t* + 1 using the following rules (Fig. 1):

<u>Rule 1</u>. Count the number of live cells *ℓ*(*t*) at time *t* in the Moore neighborhood of each *cell* in the array (i.e., 0 ≤ *ℓ* ≤ 8)

<u>Rule 2</u>. If the *current state* (i.e., state at time *t*) of the cell in question is 0 (dead) and *ℓ*(*t*) = 3, then the *next state* (i.e., state at time *t* + 1) of this cell changes to 1 (live), otherwise the *next state* remains 0

<u>Rule 3</u>. If the *current state* of the cell in question is 1 and 2 ≤ *ℓ*(*t*) ≤ 3, then the *next state* of this cell remains 1, otherwise the *next state* of this cell changes to 0

A quasi-biological narrative for these rules are: some nurturing by neighbors is needed to bring lifeless cells to life, but excessive crowding of live cells result in their death.

#### Emergent properties

Extensive simulation over the past half-century and intensive analysis of Conway’s GoL, both of which are discussed in various chapters of Adamatzky (2010) edited volume, indicate that only a limited set of values for *ℓ*, other than those laid out in Rules 2 and 3 above, define GoLs that exhibit interesting behavior. Rules 2 and 3 are the ones that define Conway’s GoL and they appear to provide the right balance between cooperative and crowding effects to produce the following veritable zoo of spatio-temporal structures:

<u>Still life motifs.</u> These are small patterns that never change (i.e., fixed over time).

<u>Blinkers.</u> These are motifs (i.e., small patterns) that switch between two configurations: i.e., they are period 2 patterns.

<u>Oscillators.</u> These are motifs that cycle among *n >* 2 sequential patterns

<u>Gliders.</u> Gliders are oscillators that move. They go through a cycle of consecutive patterns and at the start of the next cycle, the same pattern appears but has moved either vertically, horizontally, or diagonally with respect to the location of the pattern at the start of the cycle (for a rendition of Gosper’s glider see the top strip of Fig. 2)

**Figure 2.**
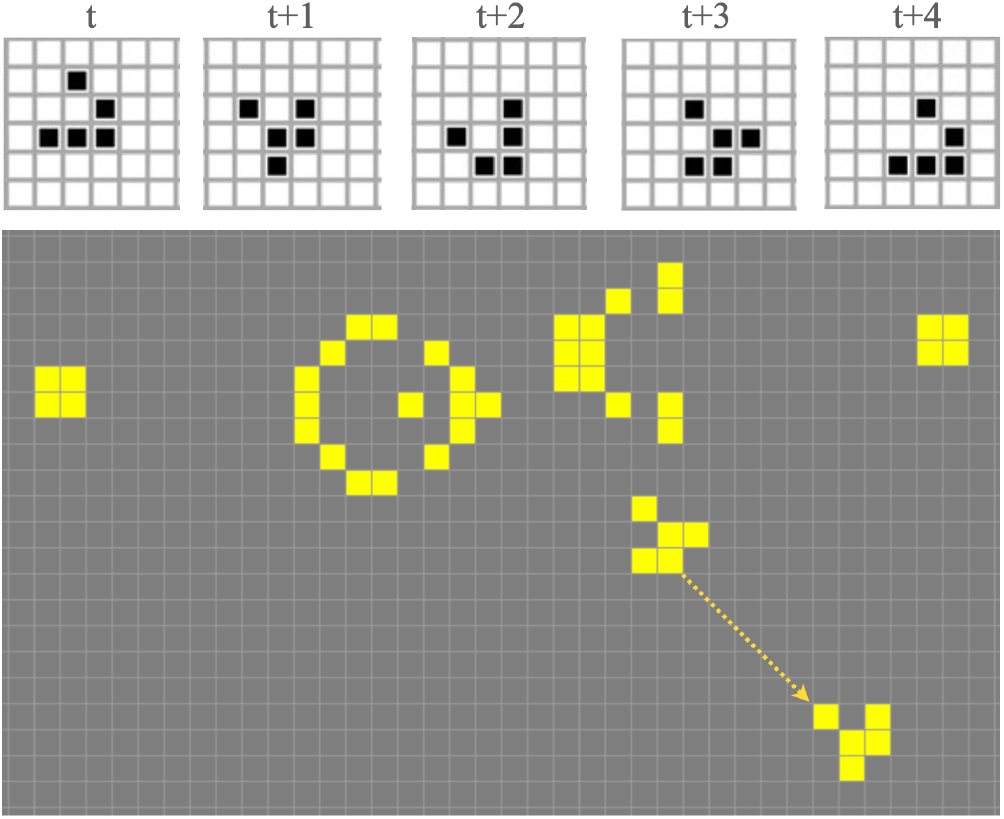
The sequence of cells at the top represents the progression of the Gosper glider over 5 times steps. Note that over this period of time we get back to the original pentagram configuration, but it has now moved one row to the left and one row down. Thus the glider moves one diagonal step in the array every five time steps. The gray and yellow cellular array below illustrates one point in the Gosper gun cycle, which launches a new glider every 30 time steps: i.e., the yellow broken arrow indicates the progression of a recently launched glider (at start of the arrow) to the previously launched glider 30 time steps earlier (at the end of the arrow).

<u>Gosper’s glider gun.</u> This is a complex 30-period oscillator (*n* = 30) that generates a 5-cell glider every 30 times steps, where these gliders move diagonally away from the oscillating gun (Fig. 2)

<u>Replicators.</u> These are patterns that after a period of time have duplicated themselves. This duplication continues over time, though some of the duplicated patterns may interfere with one another as they endeavor to occupy overlapping spaces (as discussed below in the context of one of our 4-state extensions to the game of life)

### Extensions to GoL

The particular Conway GoL presented in the previous section has the numerical designation 3/2,3 (see Table 1 for a more general discussion of this notation). This refers respectively to the conditions for a cell to come alive (*ℓ* = 3) and the bounds on *ℓ* for a cell to survive (i.e., 2 ≥ *ℓ* ≥ 3). Other configurations a such as 3,8/2,3 (i.e. a dead cell comes alive when *ℓ* ≥ 3 since the maximum possible value of *ℓ* is 3) exhibit some of the interesting properties of the 3/2,3 version (Adamatzky, 2010).

**Table 1.**
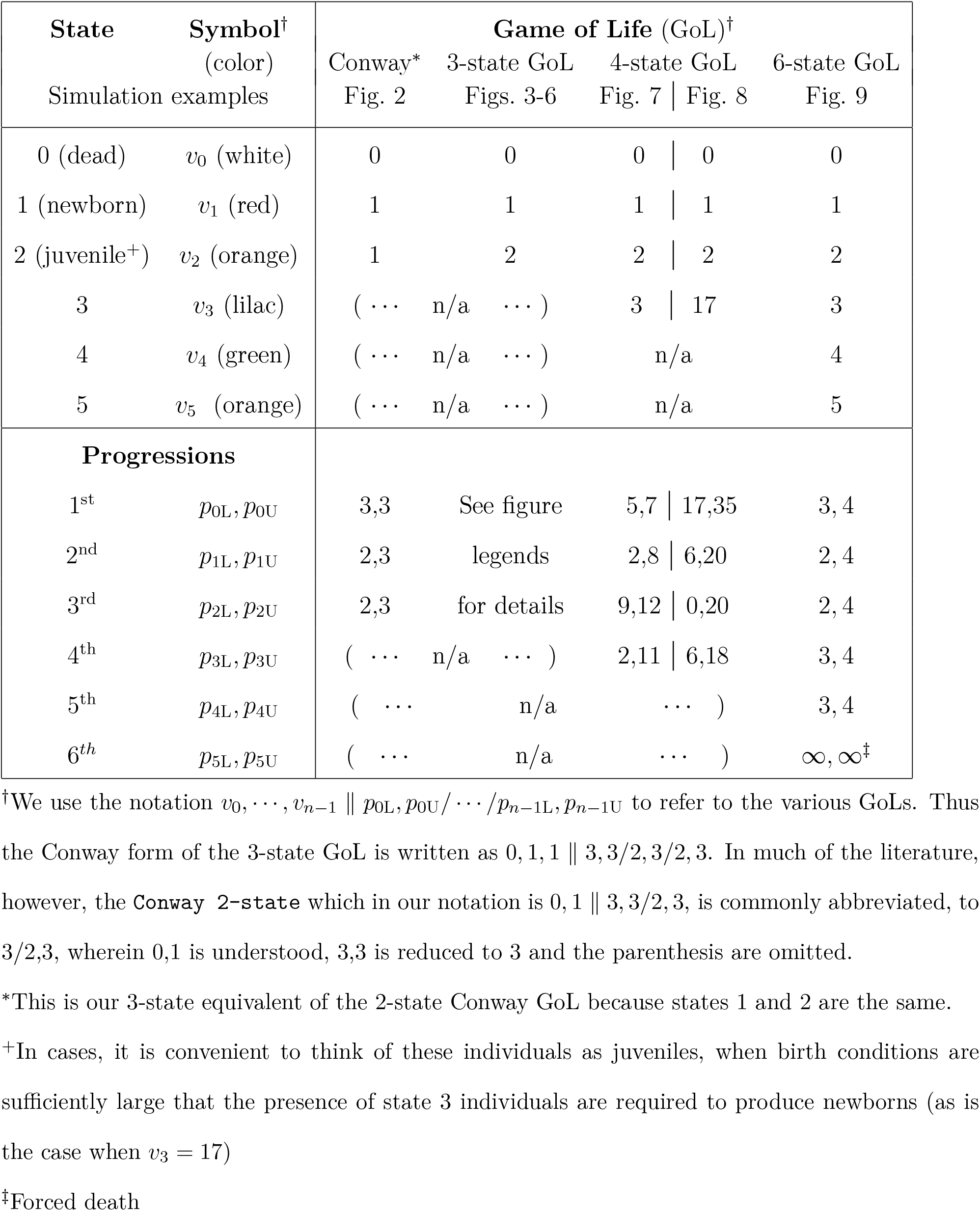
State and progression parameter values for the 3, 4-state, and 6-age Games of Life presented in our simulations

GoLs can be extended beyond these 2-state dead/live formulations to *n*-state formulations where *n* ≥ 3 (Table 1). The primary reason for moving beyond 2-state formulations is didactic rather than computational—since, as already mentioned, Conway’s GoL has the computational power of a Universal Turning Machine (Berlekamp et al., 2004). Our extended formulations provide a richer set of opportunities for student experimentation through simulation in ways that will help develop a students imagination and intuition for the complexities of discrete dynamical systems. Further, instead of students having to compete with one another to discover interesting phenomena using Conway’s already heavily explored GoL, they can more readily fire up their imaginations with the myriad more possibilities provided by exploring the extended GoLs presented here. Most importantly, as well, the RAMP applications platforms that we described in this paper can be easily used by students to carry out these explorations.

Various approaches can be taken to extending 2-state GoLs. The simplest of these is to formulate a 3-state GoL with states 0, 1 and 2 and rules for the transition of cells from 0 to 1, 1 to 2, and 2 back to 0 in terms of the neighborhood sums *ℓ*. Obviously, a more general approach would allow transitions from state 0 to 2, 1 to 0, and 2 to 1 as well. In the spirit of keeping a quasi-biological narrative to our extended game, we only consider: i) progressive transitions as the cell ages—i.e., 0 to 1 and 1 to 2 but not 2 to 1; ii) transitions that lead to death—i.e., 1 to 0 and 2 to 0. For even greater generality, we will formulate our extension in terms of *n* cell states, the first of which always has the value 0 and is regarded as the dead or lifeless state. Thus any cell can go from state *i* to state *i* + 1 for *i* = 0, …, *n* − 1 (*n* progressive transitions) or from state *i i* = 1, …, *n* − 1 back to state 0 (*n* − 1 death transitions). Of course, cellular automata “games” that have rules for transitions from one state to any other state in the context of *n* state formulations can be formulated. These, however, no longer fall under the rubrick of “Games of Life,” if GoLs are restricted to unidirectional stage transitions that model aging over time (albeit physiological and not just chronological age; see Birt et al. (2009)).

In demonstrating some of the interesting spation-temporal patterns associated with our formulation of an *n*-state GoL, we will first, consider the case *n* = 3 and then *n* = 4 (Table 1). Finally, we will discuss how to set up a *n*-state GoL when only states 3 to *n* are considered adults and at least one adult is needed in the neighbohood of a lifeless cell to bring it to life. In this *n*-state game, we will also impose the condition that progressive transitions are mandatory—i.e, an individual in state *i, i* = 1, …, *n* − 2 must in the next time interval either make the transition to the next state *i* + 1 or die, as well as imposing the finality condition that individuals in state *n* − 1 can only die in the next time interval. This provision identifies states with a notion of “age” and implies that individuals cannot live for more the *n* time periods.

The purpose of our extensions and software implementation is to provide students with applications programs that they may use to explore the behavior of these GoLs in their respective parameter spaces, thereby stimulating both their quantitative and aesthetic imaginations. As we also discuss, our 3-state GoL can be reduced Conway’s 2-state GoL and, in similar manner, our 4-state GoL can be reduced to 3-state or even 2-state GoLs.

### Our Extended GoL Formulations

Our *n*-state GoL is formulated in terms of a general set of parameter values, thereby allowing users of our RAMP implementations some freedom to explore the associated state values and transition constants parameter space. The states and the symbols used to represent these values are:

<u>Dead/lifeless cell.</u> State 0, value *v*_0_ = 0.

<u>Newborn cell.</u> State 1, and we set *v*_1_ = 1 (this is not restrictive since all values can be scaled relative to *v*_1_ without affecting the game).

<u>Living cells.</u> States *σ, σ* = 2, …, *n* − 1, *v*_*σ*_ *>* 0.

To set up our rules of the game, we first define the variable *V*_*i,j*_(*t*) to be the sum of the state values of all the cells in the Moore neighborhood of cell (*i, j*), which thus satisfies the equation

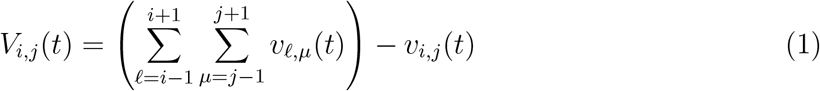

Note that since the above sum includes the value of cell (*i, j*) itself, we need to subtract this self value out from the sum.

To complete the specification of our 3-state GoL, we specify the rules the control the transition of the cells from one state to another. Here for the sake of completeness, we formulate these rules in terms of an *n*-state GoL, with cases 3 and 4 obtained by setting *n* = 3 and *n* = 4 respectively in Eq. 1 and Eqs. 2 below. For all cells *i* = 1, …, *r, j* = 1, …, *c* and time steps *t* = 0, 1, 2, … these transition rules for states *σ* = 0, …, *n* − 1 are (also see Table 1):

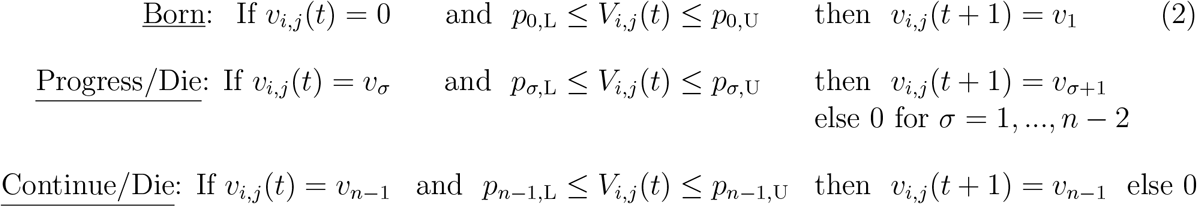

Note that when we set *n* = 3, our simulation reduces the resulting 3-state GoL to a 2-state GoL if we set (parenthetical values correspond to Conway’s GoL) *v*_1_ = *v*_2_ (= 1) and we capture Conway’s GoL as a special case when we set *p*_1,L_ = *p*_2,L_ (= 2), and *p*_1,U_ = *p*_2,U_ (=3) Also, because of the way we have written our code, once we have set *v*_2_ = *v*_1_ in our 3-State GoL RAMP, then specification of *p*_2,L_ and *p*_2,U_ is actually ignored.

### Representing and colorizing extended GoLs

A short hand notation has been developed to represent 2-state GoLs with different parameter values. We now extend this notation to represent any *n*-state GoL that is played on a rectangular array and expressed in terms of integer state values and parameter intervals controlling progression/die events. Specifically, with reference to the notation presented in the previous section, our *n*-state GoLs can be represented by the following string of integers, commas, lines and slashes *v*_0_, *v*_1_, …, *v*_*n−*1_ ∥ *p*_0L_, *p*_0U_*/* … */p*_*n−*1L_, *p*_*n−*1U_. In this notation, Conway’s 2-state game is represented by 0, 1 ∥ 3, 3*/*2, 3, which in the current short-hand notation with written as 3/2,3. In this abbreviated notation, the string 0,1 is understood and 3,3 is reduced to 3. Similarly, in our notation, if *p*_*i*L_ = *p*_*i*U_ then no information is lost by placing the single value *p*_*i*L_ rather than the order pair of values *p*_*i*L_, *p*_*i*U_ in the string. However, we will avoid this practice and, for clarity, and use the full string representation throughout.

As mentioned, implementations of Conway’s GoL typically associate the colors white/black with the states dead/live, or in some cases, gray/yellow, as seen in Figs. 1 and 2. The *n*-state GoL can be visualized on a cellular array using shades of gray or, more dramatically, distinct colors for each state. We use white, red, and orange as the default colors for states 0, 1, and 2 in our 3-state GoL, with the addition of lilac to represent state 3 in our 4-state GoL (Table 1). These color defaults can be overridden by the user in our GoL RAMP implementations.

### Initial configurations, motifs, and phases

The behavior our extended GoL can roughly be thought of as having two possible distinct phases: an initial phase heavily influenced by the starting configuration (aka initial conditions) of the simulation and a final phase that reflects the dynamical properties of the system rather than a particular set of initial conditions. This statement excludes those initial conditions set up to generate specific initial configurations—which we well call motifs—such as the still-life, blinker, oscillator, glider and replicator motifs discussed above in the context of Conway’s GoL.

One way to identify this possible zoo of motifs is to keep rerunning simulations on a toroidal array, starting from randomly configured initial conditions and searching for the emergence of such motifs before they get gobbled up by colliding with other motifs. Once such putative motifs have been identified, they can be isolated and run from set-piece initial conditions configured (as we shall illustrate in some of our selected examples) to identify the temporal sequences of these motifs in the absence of surrounding elements that may lead to their demise. These set piece simulations can be carried out on arrays with either edge or wrap boundary conditions (which can be selected on the RAMP).

Additionally, large scale regularized initial conditions, can lead to the production of cyclic patterns at the scale of the full array itself when simulations are executed on arrays with either edge or wrap boundary conditions. Below we provide a guide to setting up such regular arrays, along with mathematical descriptions for setting them up, as well as setting up random and motif-specific initial configurations. In the RAMP itself, specific RAMs are available for implementing regular and random initial configurations with the freedom to select parameter values that control elements of these configurations.

<u>Regular initial conditions.</u> Regular initial conditions can be set up using modular arithmetic. We do this by defining a variable *ℓ*_*i,j*_ = (*i* + *j*) mod *r* (i.e., the reminder we get when (*i* + *j*) is divided by *r*). We then set up a list of *r* ordered pairs that specifies what the state of cell (*i, j*) (second entry in the order pair) will be when *ℓ*(*i, j*) = 0, …, *r* − 1 (first entry in the ordered pair). For example, in the 4-state game used to set the initial conditions seen in Fig. 7, the lists for the four panels with *r* = 6, 7, 8, and 9 respectively (going from left to right) are: {(*ℓ*_*i,j*_ = 0, `state` = 1), (*ℓ*_*i,j*_ = 1, `state` = 2), and (*ℓ*_*i,j*_ = *q*, `state` = 0) for *q* = 2, ‥, mod *r*}. When the initial configuration RAM is selected for the start of a simulation, the user will be able to specific the values of *r*, as well as the states associated with each of the reminders when (*i* + *j*) is divided by *r*: i.e., the regular specification is

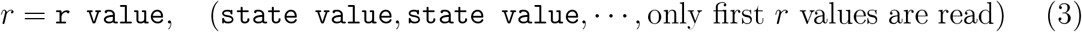

<u>Random initial conditions.</u> Random initial conditions are easily set up using the computers random number generator. For example, for each cell we can draw a cell-specific random value *z*_*i,j*_ ∈ [0, 1] (every value between 0 and 1 is equally likely). Then an *n*-state game, we specific *n* − 1 parameter values *z*_*ν*_, *ν* = 1, …, *n* − 1 assign cell (*i, j*) to one of the *n* states as follows

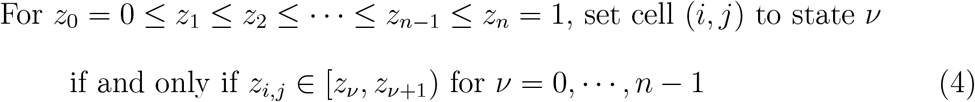

In this case the expected proportions of cells that are initially in state *ν* respectively is *z*_*ν*_ − *z*_*ν*+1_, *ν* = 0, …, *n* − 1. Note, if *z*_*ν*_ = *z*_*ν*+1_ then clearly the interval [*z*_*ν*_, *z*_*ν*+1_) contains no points.

<u>Motif-specific initial conditions.</u> In our case, we set motif-specific initial conditions by first setting all cells to 0 (i.e., a `lifeless array`) and then using our mouse to toggle selected cells to particular states by first pressing the command button 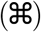 and then clicking the mouse one or more times while pointing to the cell being set. Specifically, one click turns the cell to state 1, two clicks to state 2, and so on upto *n* clicks for the final state *n*, with *n* + 1 clicks returning to state 0.

Apart from the set-piece initial configurations—which we may set up to illustrate particular still-life, oscillatory, glider, and replicator motif sequences—simulations that start from regular or regular initial configurations, or set piece configurations that are not part of a particular motif sequence, have two distinct phases: a burn in phase that steers the system towards a final phase in which: i.) all cells are either dead, ii) the array supports a sparse set of non-interfering still-life, oscillatory or glider motifs, or iii) the array-wide pattern emerges that has regions exhibiting oscillatory behavior where the period of oscillation for the whole pattern itself may be so vast as not to be detectable. These ideas will be clarified, as we present specific examples of simulations starting from regular, random, or set-piece initial configurations.

### 3-State GoL Examples

In this section we provide examples using all three types of initial configurations in the context of 3-state GoLs. We begin by exploring the simplest possible extension of Conway’s GoL to 3 states: we set *v*_2_ = 2 and keep all the other parameters the same (i.e., the GoL 0, 1, 2 ∥ 3, 3*/*2, 3*/*2, 3). In this case, all random starting positions converge on configurations typical containing two still-life motifs, as well possibly one or more of three glider motifs (Fig. 3). The still-life motifs are two live cells side by side (vertically or horizontally) and live cells that are diagonal neighbors. The glider motifs are two sequences of patterns of period 5 with patterns and one of periods 6 (Fig. 3). These are not necessarily the only gliders associated with 0, 1, 2 ∥ 3, 3*/*2, 3*/*2, 3, but if others are to be found, they arise much less frequently from random starting conditions than the two 5-period gliders depicted in Fig. 3, and at least somewhat less frequently than the 6-period glider.

**Figure 3.**
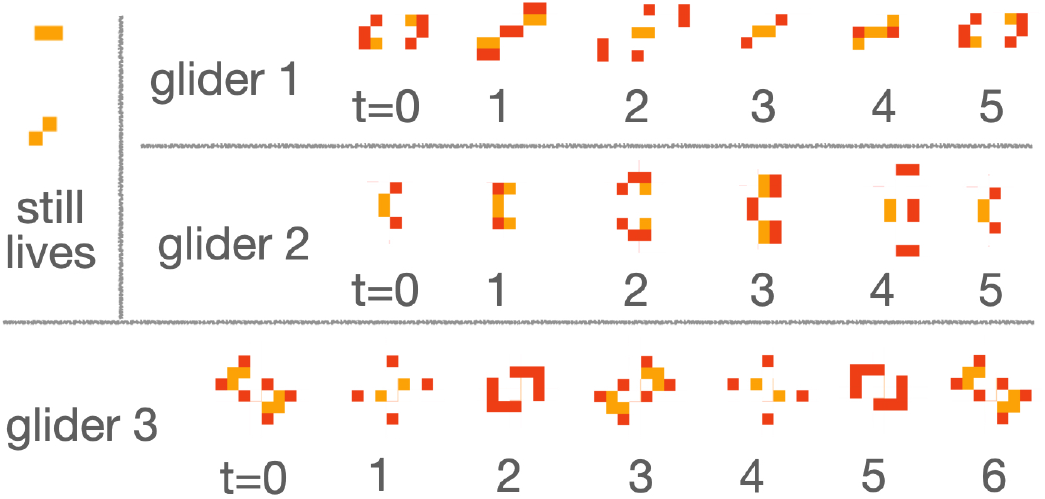
Still lives and gliders produced by a one state addition with a one unit increment in value to the Conway GoL, but with the transition parameters remaining the same: i.e. the basic 3-state GoL 0, 1, 2 ∥ 3, 3/2, 3/2, 3 (newborn and live individuals in red and orange respectively). The still lives are two adults side-by-side either vertically, horizontally, or diagonally (two shown plus two obtained from 180 degree rotations). Three known gliders are: two period 5 and one period 6, where the latter consists of a period 3 series followed by its 3 mirror images.

By setting up a GoL in which the largest state value had been increased from 1 at 2, the neighborhood values *V*_*i,j*_(*t*) (Eq. 1) can now be twice as large as in Conway’s 2-state GoL. Thus, it seemed reasonable to explore the behavior of our 3-state GoL by increasing the GoL birth parameters from Conway’s 3,3 to 3,4 and tweaking the newborn progression range to 2,4, but keeping the density at which the live state would still persist at 2,3. Thus, we explored the behavior of the GoL 0, 1, 2 ∥ 3, 4*/*2, 4*/*2, 3, starting from a series of random initial conditions. In these simulations, we were able to putatively identify two different gliders, both more complex than the three gliders that we found in the GoL 0, 1, 2 ∥ 3, 3*/*2, 3*/*2, 3 (Fig. 4). The first of these (Glider 1, Fig. 4) is an 8-period oscillator that has an obvious diagonal progression moving 4 squares every 8 steps. The 8 patterns associated with this glider are all diagonally symmetric with the simplest of these involving `2 newborn + 6 live` and the most complex involving `6 newborn + 12 live`. The second glider (Glider 2, Fig. 4) is quite astonishing. It has a 46 period cycle (23 patterns followed by their 90 degree clockwise rotated mirror images) and lacks diagonal or other symmetries. This lack of symmetry gives the visual impression that this glider oscillates back and forth along one diagonal axis while steadily progressing along the other diagonal axis. The simplest of the 23 patterns involves `4 newborn + 3 live` while the more complex of the 23 patterns involves 9-11 newborns and 9-12 live individuals.

**Figure 4.**
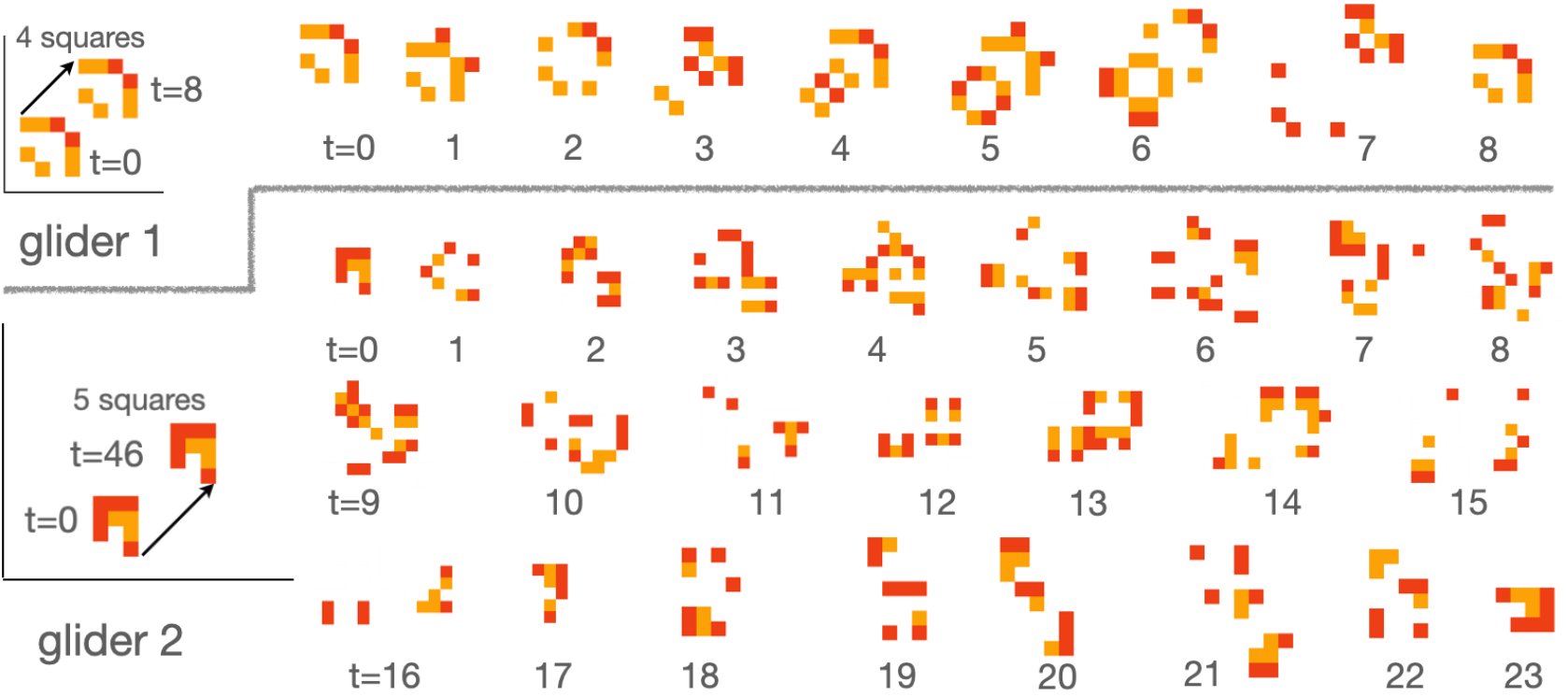
Two known gliders of the basic 3-state GoL 0, 1, 2 ∥ 3, 3/2, 4/2, 3 (newborn and live individuals in red and orange respectively). Glider 1, period 8, is a sequence of 8 diagonally symmetric patterns that moves the starting pattern 4 squares along the diagonal every 8 steps. Glider 2, period 46, is a sequence of 23 patterns followed by 23 of their mirror images rotate clockwise by 90 degrees, and moves the starting pattern 5 squares along the diagonal every 46 times steps.

**Figure 5.**
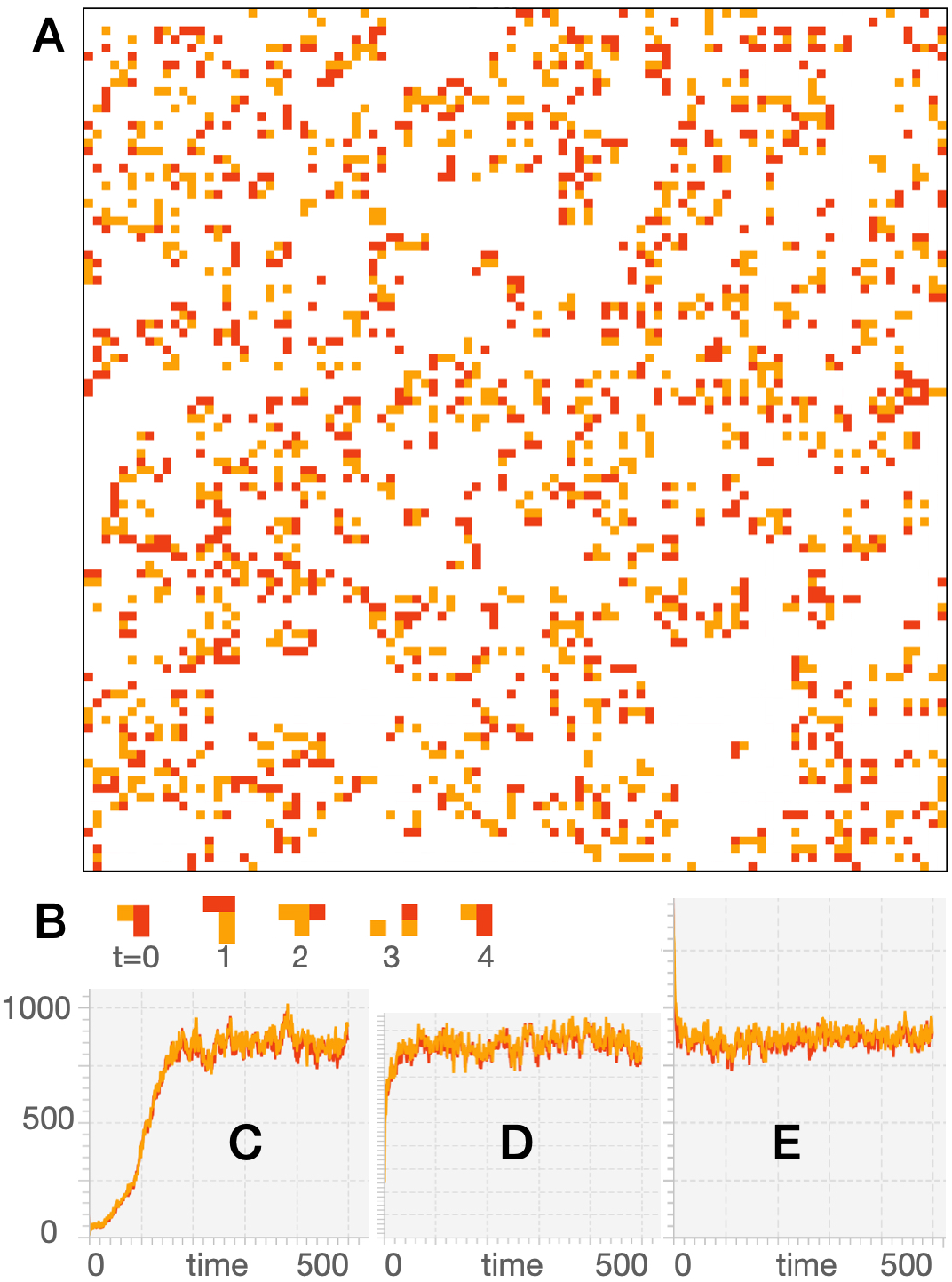
A. From a random initial configuration, irrespective of the initial conditions, provided they are not too sparse (i.e., the array dies before it can get going), simulations of the GoL 0, 1, 2 ∥ 3, 3/2, 4/3, 4 on a 100×100 toroidal array settles on a seemingly every changing chaotic pattern (which will eventually repeat itself, as discussed in the text) that varies between approximately `8-9% newborn` (red) and a comparable number of `live` (orange) cells (white cells are `dead`). **B**. This GoL, produces at least one type of glider, but such gliders appear fleetingly as they emerge from the chaos and then are consumed by colliding with other newborn and live cells. The oscillatory band of between 800-900 newborn and a similar number of live cells approached within the following period from the given randomly assigned starting conditions: **C**. approximately 150 steps, 1% of each cell type; **D**. approximately 60 steps, 5% of each cell type; **E**. approximately 10 steps, 15% of each cell type. The outcome remains the same even with the starting number of newborn and live cells vary by a factor of 10 or more.

We next considered the behavior of the GoL used to produce the gliders in Fig. 4, except we increased the live persistence conditions from 2,3 to 3,4. Simulations of this GoL on a toroidal 100 × 100 array from random starting conditions converged on a seemingly chaotically changing pattern of newborn and live cells with comparable numbers of both, oscillating in a band between 800-900 cells (i.e., a little over 80% of the cells at any step beyond the initial burn-in phase were dead). This band indicated no periodic behavior, though we note that being a finite state system, our toroidal array has 3^1^0, 000 (10^12^) states (ten thousand cells, each of which can be in one of three states), so at some point one of the states must reoccur. Being deterministic, when a state reoccurs, then all the succeeding states are repeat and hence an oscillatory situation has set in. However, 3^1^0, 000 is such a vast number that the periodicity of this simulation may never be found, even if we keep computing until our solar system ceases to exist. We did, however, identify one putative glider from these random simulations, which we then checked and found that it was in fact a glider by simulating the system from any one of the 4 motif-specific initial conditions shown to be part of the 4 period glider sequence in Fig. 4B.

We then moved on to exploring the behavior of a 3-state basic GoL 0, 1, 2 ∥ 3, 4*/*3, 4*/*2, 5 that, compared with our previous 0, 1, 2 ∥ 3, 4*/*2, 4*/*2, 3 GoL, had tighter conditions for the generation of newborns but an increased neighborhood value range at which live cells would persist. We simulated this GoL on a 100 × 100 bounded rectangular array (i.e., edge boundary conditions), starting from a diagonally banded initial configuration consisting of consecutive diagonal bands of newborns (red cells) and live individuals (orange cells) separated by 4 lifeless diagonals (dead cells are white) in between (i.e., in Eq. 3 we specify *r* = 6, (1, 2, 0, 0, 0, 0, …)). The state of the array is depicted, as labeled at times *t* = 0, 4, 41, and 100 (the four smaller rectangles on the left in Fig. 6). The simulation converges to a pattern that repeats itself every 11550 (= 2 × 3 × 5 × 5 × 7 × 11) steps. The large rectangle shows one of the points in this 11550 period cycle. The smaller identified rectangular sections on the right contain patterns that repeat themselves every (from top to bottom) 150, 10, 30, and 66 time steps.

**Figure 6.**
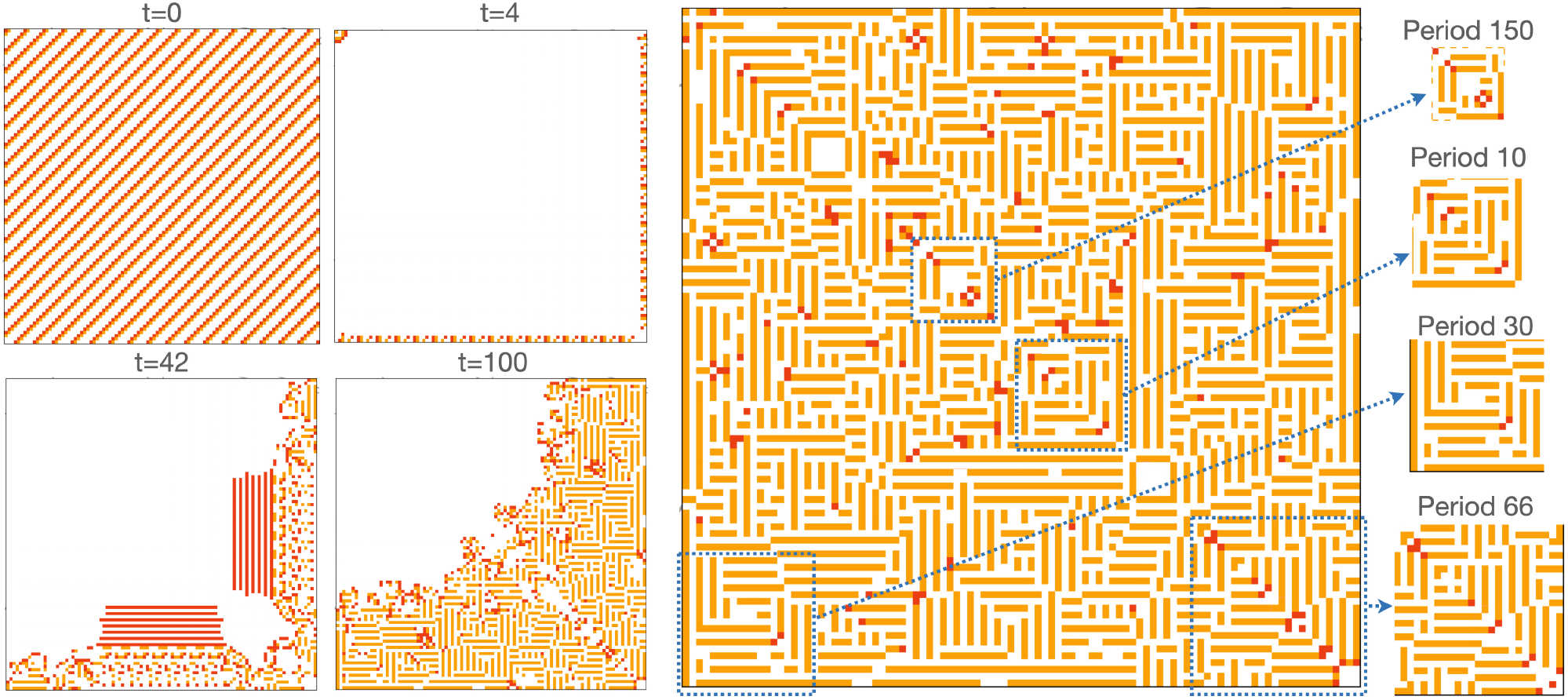
A simulation of the 3-state basic GoL 0, 1, 2 ∥ 3, 4/3, 4/2, 5 (Table 1) on a recangular array (i.e., edge boundary conditions) with r = 100 rows and c = 100 columns (newborn and live individuals in red and orange respectively) with regular initial condition r = 6 and array specification (1, 2, 0, 0, 0, 0) (see Eq. 3). The four smaller rectangles on the left show that state of the array at times t = 0, 4, 42 and 100. The large rectangle to the right of these 4 represents one of the states that is part of an oscillatory cycle, which repeats itself every 11550 (= 2 × 2 × 3 × 5 × 7 × 5 × 11) steps. The rectangular subregions identified by blue dotted lines are extracted on the right-hand-side with the period of the elements inside these rectangles labeled above them.

The initial conditions we set up are just at the edge of being sufficient for colonization to begin on two of the four boundaries, while wiping out all live cells on the remaining two of the boundary conditions and in the interior of the (*t* = 4, Fig. 6). These boundary populations started expanding, keeping the diagonal symmetry imposed from the start. For example, we see in Fig. 6 at time *t* = 42 that the interior region is being invaded by a front of newborn cells. This front leaves behind domains of live horizontally or vertically striped cells with unsettled boundaries between domains support individual or small groups of newborn cells (e.g., see *t* = 100 rectangle in Fig. 6). Finally, the simulation settles into its periodic behavior with different areas of the whole array exhibiting regions with faster periods that are factors of the whole array period. Interestingly, a relatively small array on the diagonal just above the center has period 150, while a much larger area in the lower left corner of the array has period 30. The periods of any one of these subregions are necessarily multiples of the factors of 11550, such as 66 (2 × 3 × 11) or 30 (2 × 3 × 5). Once the simulation gets going it soon settles into some pattern that has the same basic theme of horizontal and vertical stripped domains of dead/live cells with domain boundaries supporting oscillating patterns of newborn cells. The diagonal symmetry of the pattern that we see in Fig. 6 only occurs when the array is square and the initial conditions are diagonally symmetric themselves.

### 4-State GoL Examples

The most natural way to extend the 3-state GoL to a 4 state GoL is to add an additional state of value 4 to obtain the first of the 4-state GoLs listed in Table 1. One can then play around with different sets of progression values until some interesting results emerge. Some 4-state GoLs will have parameter values that are too liberal or too restrictive too produce interesting final phase results. However, many different parameter combinations of the 4-state GoL 0, 1, 2, 3 ∥ *p*_0L_, *p*_0U_*/p*_1L_, *p*_1U_*/p*_2L_, *p*_2U_*/p*_3L_, *p*_3U_ are likely to produce all kinds of interesting and, often, surprising results. Here, by way of illustration, we present a couple of examples, with the vast field of possibilities of different parameter values, initial configurations, and array boundary conditions left to students and researchers interested in cellular automata dynamics to discover for themselves.

Our first example pertains to the 0, 1, 2, 3 ∥ 5, 7*/*2, 8*/*9, 12*/*2, 11 GoL. We simulated this GoL on a 100 × 100 toroidal array from a one parameter set of regular initial starting conditions defined using the variable *ℓ*_*i,j*_ = (*i* + *j*) mod *r*. For values of *r* ranging from 3 to 12, we specified that cell (*i, j*) starts in state 2 if *ℓ*_*i,j*_ = 0, starts in state 3 if *ℓ*_*i,j*_ = 0, and otherwise starts in state 0 for all other values of *ℓ*_*i,j*_ = 0. These simulations only produced results for *r* = 6, 7, 8 and 9 (Fig. 7). For *r <* 6 and *r >* 9 the simulations ended up in just a few time steps with an array dead cells because conditions with either initially too crowded (*r* = 3 to 5) or too sparse (*r >* 9) to sustain interesting life. The simulations obtained in the remaining 4 cases are illustrated in Fig. 7

**Figure 7.**
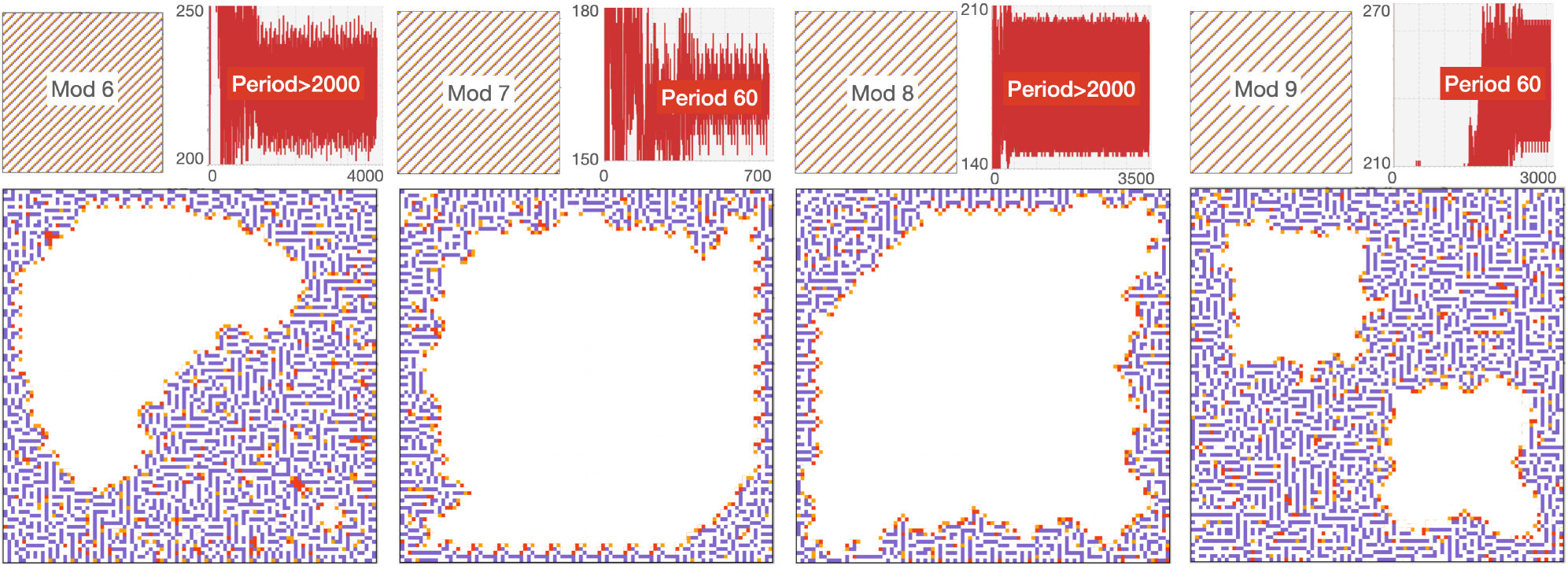
A simulation of the 4-state basic GoL 0, 1, 2, 3 ∥ 5, 7/2, 8/9, 12/2, 11 (Table 1) on a torus with r = 100 rows and c = 100 columns. Newborn (v_1_ = 1), juvenile (v_2_ = 2) and adult (v_3_ = 3) individuals are respectively colored red, orange, and lilac. The four cases, from left to right respectively, represent starting conditions in which a diagonal of juvenile individuals is immediately followed by a diagonal of adults, followed by 4, 5, 6 or 7 consecutive diagonals of dead (v_0_ = 0) individuals, depending on the value of r = 6, 7, 8, 9 chosen to generate the starting pattern using the function ℓ_i,j_ = (i + j) mod r, as described in the text regarding that setting up of regular initial conditions. E.g., in Eq. 3 we set our values as r = selected value, (2, 3, 0, 0, 0, 0, 0, …).

Our second examples is inspired by the biological consideration of regarding the zero state as a lifeless/dead cell with *v*_0_ = 0, the first non-zero state as newborn with *v*_1_ = 1, the second non-zero state as juvenile with *v*_2_ = 2, and the third non-zero state as adult with its value *v*_3_ to be selected. If the third non-zero state is an adult and we require at least one adult for a lifeless cell to come to life, then a lifeless cell surrounded by 8 juveniles—which implies *V* = 16 for the cell in question (Eq. 1)—implies that we should set *v*_2_ ≥ 17 to avoid the event of 8 juveniles raising one newborn without an adult present. If we set *v*_3_ = 17, then we are interested in the GoL

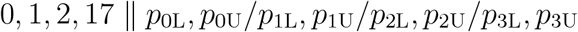

We could, of course, consider the situation in which each adult requires at least one helper to raise a newborn, in which case, we would need to set *v*_3_ = 18. We will leave this latter case to the reader to explore.

In the former case, we explored the GoL 0, 1, 2, 17 ∥ 17, 35*/*6, 20*/*0, 20*/*6, 18. In particular, we undertook a search for oscillators by simulating this GoL from a relatively sparse random initial conditions (93% dead, 4% newborn, 1% juvenile, 1% adult). For this game we found both a glider and a replicator, as illustrated in Fig. 8.

**Figure 8.**
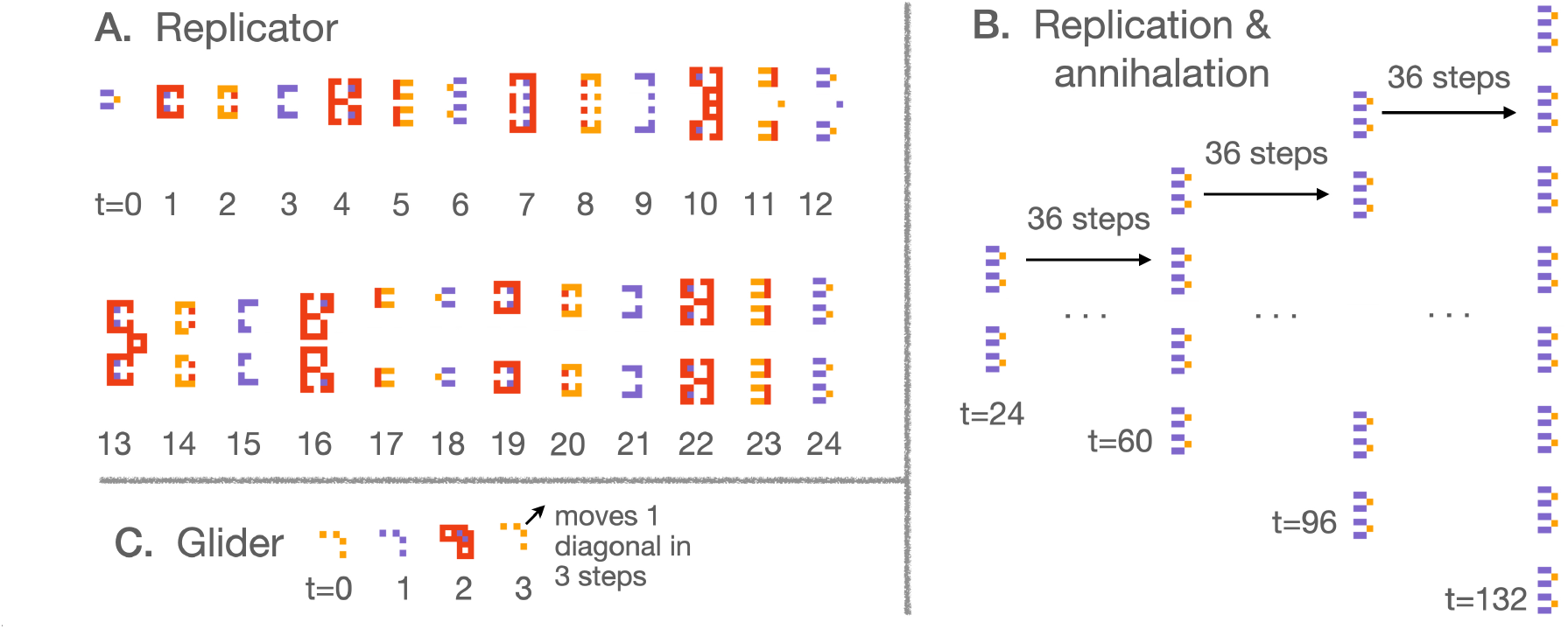
Associated with the 4-state GoL 0, 1, 2, 17 ∥ 17, 35/6, 20/0, 20/6, 18 (newborn, juvenile, and adult individuals in red, orange, and lilac respectively) is **A**. A motif that replicates itself in mirror image form after 6 steps, and shows a four-fold increase after 24 steps. **B**. The replication process, however, is not a simple doubling every so many steps because replication motifs interfere with one another along the axis perpendicular to the symmetry axis of the motif, with internal pairs of motifs along this axis appearing and disappearing over time as the replication process expands in both directions along the said perpendicular axis. **C**. We also discovered a simple period 3 glider that moves along a diagonal one square in every 3-step cycle.

### 6-State Age-specific GoL Examples

As our final demonstration, we consider a 6-state GoL in which all individuals die after being in state 6 for one step. We could, of course, had the same final state condition for one of our 4-state GoLs, and only include this illustration for emphasizing the open endedness of our formulation. We do not, however, provide a RAMP to implement 6-state GoLs, but instead leave it to the reader to build their own versions, perhaps using Numerus Designer^®^ to do so.

The simplest case of a 6-state GoL with forced termination of the last state is

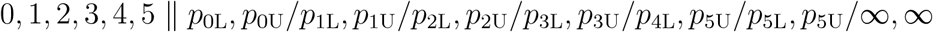

(we note the final limits could just as well be 41,41 instead of ∞, ∞ becasue 40 is the largest value that *V* can be for the neighborhood of any cell—see Eq. 1). The particular case of this class of GoLs that we explore is one in which all individuals die, no matter their state, whenever neighborhood values *V* exceed 4. In addition, we set the lower bounds for births at 3 and for transition of newborns (state 1) to state 2 and state 2 cells to state 3 is 2. Finally, the lower bounds for state 3 to state 4 and state 4 to 5 where set at 0. Thus resulting GoL we explored is: 0, 1, 2, 3, 4, 5 ∥ 3, 4*/*2, 4*/*2, 4*/*3, 4*/*3, 4*/*∞, ∞ (Table 1).

This GoL exhibits an exceedingly rich behavior that we can only touch on here (Fig. 9). Beyond the various motifs that it may produce (which we have not yet tried to identify)—still lives, oscillators, gliders, and replicators—its final phase behavior has interesting properties when starting from certain motifs that have a two or four-way symmetry (Fig. 9B). If the conditions are right for the number of cells to grow (Fig. 9B) from this initial motif (i.e., not peter out to nothing or morph into a small number of still-life, oscillator, or glider motifs) to an array-filling dynamic state that is an almost ever changing complex pattern (i.e., the period of the pattern is so vast that we are unlikely to ever see it repeated), then several centers of symmetry emerge, as depicted in Fig. 9D.

**Figure 9.**
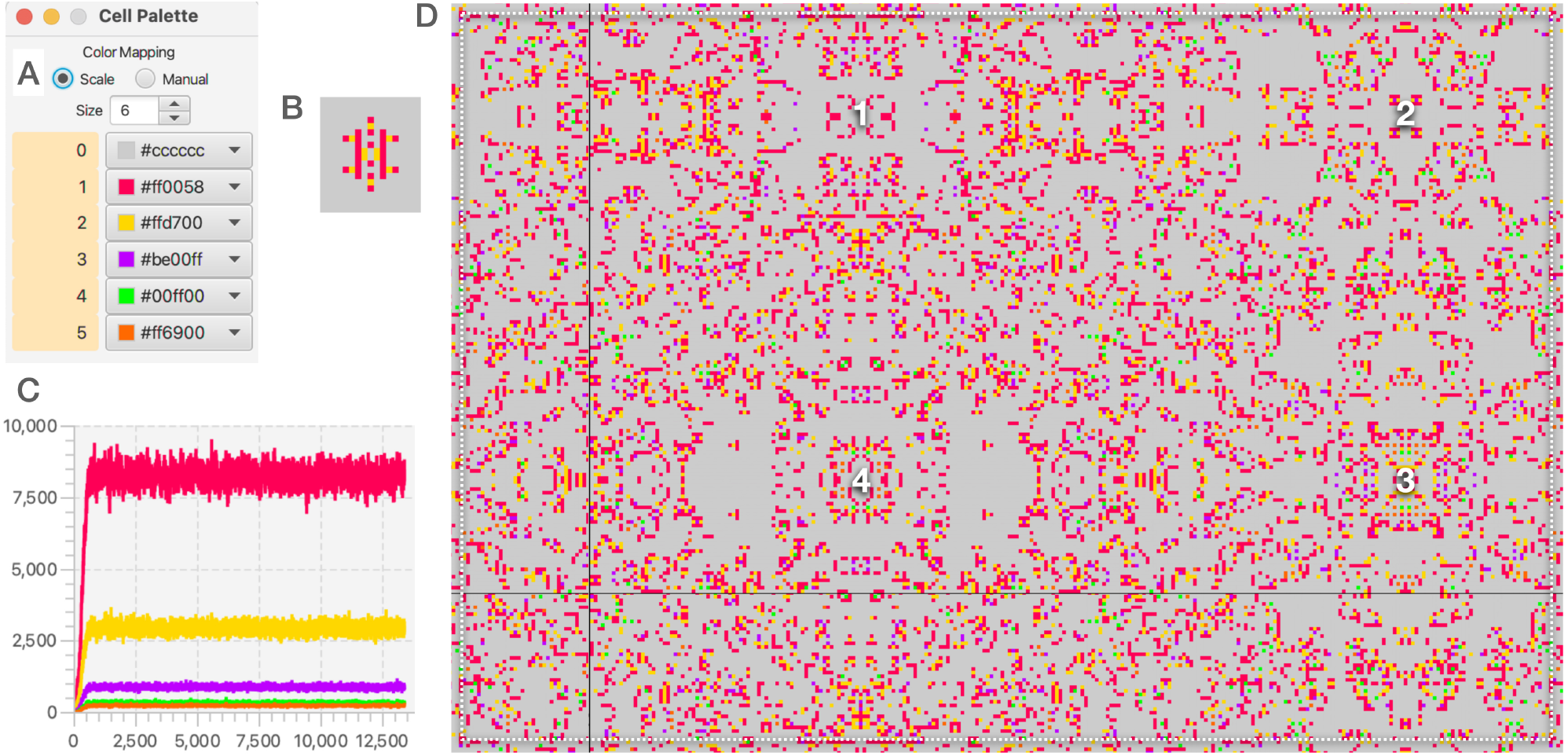
A simulation on a 200 × 300 toroidal cellular array of the 6-state GoL 0, 1, 2, 3, 4, 5 ∥ 3, 4/2, 4/2, 4/3, 4/3, 4/∞, ∞ using the color palette show in **A**. The simulation began with the initial motif **B**. This motif increases over time to completely fill the array by time t = 1000, where we see in graphs **C** that the number of cells in the five non-zero state 1-5 have leveled off to oscillate around a set of average numbers of cells. These percentages of the number of cells in each state approximately average 13.5% (state 1), 4.7%, 0.14%, 0.05% and 0.04% cells respectively for the rest of the simulation, which we terminated around t = 13, 500. The pattern in array **D** (area we have marked to be within the broken white border) is the one we captured at the end of the simulation, but it also varies over time though retaining fixed centers of symmetry throughout the rest of the simulation. In this array we have labeled 4 regional symmetry centers that persist beyond the initial simulation phase. The thin black lines correspond to the regions of the array we have patched together to depict our toroidal surface as a rectangle in such as way to make most visually obvious the extents of the regional areas of symmetry.

### RAMP Implementation

The three and four state GoLs presented here and other variations that fit within the constraints imposed by Eqs. 1 and 2 for the case *n* ≤ 4 can be implemented using our Numerus RAMPS, downloadable at our Numerus^®^ website along with our Numerus Studio applications program to implement these RAMPs. Also a pdf file can be downloaded at Numerus Studio that contains a description of how to use the current set of RAMPs downloadable at this web.

Finally, as already mentioned, many different ways of extending Conway’s GoL exist. In particular, the constraint that cells can only progress from state *i* to *i* + 1, *i* = 0, …, *n* − 2 (as implied by Eq. 2) can be changed and, for example, transitions of state *i* to other states *j* = 0, …, *n* − 1 can be determined according to particular ranges of neighborhood values *V*. RAMPs to implement such GoLs, including the 6-state GoL discussed in this paper, can be built using our Numerus Designer Applications program downloadable at the Numerus Inc Website.

## Discussion

In this paper, we have introduced the reader to Conway’s Game of Life (GoL), and to a class of extensions that, in the context of the use of our accompanying RAMPs, greatly facilitate the exploration of properties of cellular automata by curious students with no particular computational or mathematical training. Users of our RAMPs are explicitly exposed to the ideas of numerical simulations and cellular automota, while getting implicit exposure to concepts in complex systems theory. They are also get the opportunity, through exposure to our the RAMs (runtime alterable modules) that our part of our RAMP (runtime alterable model platform) construction, to alter small pieces of JAVA code.

The simulations and resulting patterns that we illustrate, suggest a number of interesting mathematical questions that can either be addressed through simulation or theoretical analyses. For example, on a toroidal cellular array of size *r* × *c*, how many gliders associated with a particular GoL (e.g., see Figs. 3-5 can coexist without interfering with one another. Also, questions arise regarding the number of symmetry centers and the dimensions of the symmetry rectangles they support in GoLs of the type considered in Fig. 9.

As we have seen in our illustrative examples, the final phase of various simulations produces patterns that can be characterized by a stationary distribution of the number of cells of each type over time and, depending on whether or not the starting pattern was regular, with associated symmetries. An additional way to characterize the final phase is to record the state of each of the individual cells in the array during the final phase and record its periodicity (fixed values have periodicity of 1). If we then look at all the cells in a local rectangle that we may identify, much as we have done in Fig. 6, then the period of the pattern that emerges in that rectangle will be the lowest common multiple of the periods of all the cells in the identified rectangle.

Finally, we emphasize that many different ways of extending Conway’s GoL exist. In particular, the constraint that cells can only progress from state *i* to *i* + 1, *i* = 0, …, *n* − 2 (as implied by Eq. 2) can be changed and, for example, transitions of state *i* to other states *j* = 0, …, *n* − 1 can be determined according to particular ranges of neighborhood values *V*. RAMPs to implement such GoLs can be built using our Numerus Designer App that can be obtained at our Numerus website.

### Concluding Remarks

The illustrations we have presented here, are the equivalent of opening a vast box of “goodies” and only picking a few of them at random to whet our appetite for generating all kinds of spatial patterns. Many of the remaining goodies may be similar to those presented here, but it is likely that many others will hold quite some surprises for the users of our RAMPs who discover them, or anyone else who sees them for the first time. We can only hope that our RAMP helps stimulate an aesthetic appreciation for the wonder and beauty of patterns produced by what at the outset seems just a simple and dull set of rules that embody the mathematical description of cellular automata.

#### Guide to using the RAMPs

A guide to using the Numerus RAMPs presented in this paper can be found at the Numerus Studio web page.

